# The origin of universal cell shape variability in a confluent epithelial monolayer

**DOI:** 10.1101/2021.08.21.457184

**Authors:** Souvik Sadhukhan, Saroj Kumar Nandi

## Abstract

Cell shape is fundamental in biology. The average cell shape can influence crucial biological functions, such as cell fate and division orientation. But cell-to-cell shape variability is often regarded as noise. In contrast, recent works reveal that shape variability in diverse epithelial monolayers follows a nearly universal distribution. However, the origin and implications of this universality are unclear. Here, assuming contractility and adhesion are crucial for cell shape, characterized via aspect ratio (AR), we develop a mean-field analytical theory for shape variability. We find that a single parameter, *α*, containing all the system-specific details, describes the probability distribution function (PDF) of AR; this leads to a universal relation between the standard deviation and the average of AR. The PDF for the scaled AR is not strictly but almost universal. The functional form is not related to jamming, contrary to common beliefs, but a consequence of a mathematical property. In addition, we obtain the scaled area distribution, described by the parameter *µ*. We show that *α* and *µ* together can distinguish the effects of changing physical conditions, such as maturation, on different system properties. The theory is verified in simulations of two distinct models of epithelial monolayers and agrees well with existing experiments. We demonstrate that in a confluent monolayer, average shape determines both the shape variability and dynamics. Our results imply the cell shape variability is inevitable, where a single parameter describes both statics and dynamics and provides a framework to analyze and compare diverse epithelial systems.

---

D’Arcy Thompson argued, in his book *On Growth and Form*, physical principles could explain tissue packing and cell shape [1]. Shape formation of tissues and organs during embryogenesis is a long-standing, fascinating problem of developmental biology. Since cells are the functional units of a tissue, shapes in the organs must originate at the cellular level [2–5]. Cell shapes are vital in both health and disease. As cancer progresses [6, 7], as asthma advances [7–10], as wounds heal [11, 12], as an embryo develops [10, 13], cells progressively change their shape. Besides, cell shape may influence crucial biological functions, such as cell growth or selective programmed cell death (apoptosis) [14], the orientation of the mitotic plane [3, 15, 16], stem cell lineage [17, 18], terminal differentiation [19, 20], and division-coupled interspersion in many mammalian epithelia [21]. Moreover, the nuclear positioning mechanism in neuroepithelia depends on cell shape variation [22]. Thompson regarded cell-to-cell shape variability as a biologically unimportant noise [1]; however, it is now known that shape variability is not an exception but a fundamental property of a confluent cellular monolayer [23]. In a seminal work, Atia *et al* [10] showed that cell shape variability, quantified by the aspect ratio (AR), follows virtually the same distribution across different epithelial systems. But, the origin of this near-universal behavior, and whether it is precisely universal, remains unclear.

Reference [10] argued that this behavior in an epithelial monolayer is related to the jamming transition. Previous works have established the similarity in dynamics between cellular monolayers and glassy systems [8, 24, 25]. The glassy dynamics in confluent systems has been argued, via vertex model (VM) simulations, to be controlled by a rigidity transition akin to the jamming transition [8, 26]. In a jammed granular system, the statistics of tessellated volume, *x*, follows a *k*-Gamma distribution, *P* (*x, k*), where *k* is a parameter. From this association, Ref. [10] conjectured the viability of describing the AR distribution via *P* (*x, k*). However, such an association has several problems: (1) The glassy dynamics in a cellular monolayer is also reasonably described by Voronoi and cellular Potts models (CPM) [27, 28]. In these models, in contrast to the VM, there is no rigidity transition [28, 29]. Hence, the relevance of jamming physics to an epithelial system remains unclear. (2) More important, the experiments and simulations in Ref. [10] show that this distribution persists even in the fluid regime, where jamming physics is not applicable even within the VM. (3) The analytical derivation of the *k*-Gamma distribution in granular packing [30] relies on the fact that *x* is additive, whereas, as the authors of Ref. [10] rightly point out, AR, *r*, is not. Thus, there *exists no rigorous basis for the applicability of the k-Gamma distribution* [10].

Yet, Ref. [10] and several subsequent works [31–34] have shown that the probability distribution function (PDF) of scaled AR, *r*_*s*_, in diverse biological and model systems is described by *P* (*r*_*s*_, *k*), and the value of *k* is almost universal, around 2.5. Furthermore, the standard deviation, *sd*, vs the mean AR, 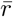, follows a universal relation [10, 33, 35]. What is the origin of this universality? What determines the value of *k* ∼ 2.5? How is the PDF of *r* related to the microscopic properties of a system? Answers to questions like these are crucial for deeper insights into the cell shape variability and unveiling the implications of the universality. However, it requires an analytical theory that is rare in this field due to the inherent complexity of the problem and the presence of many-body interactions.

In this work, we develop a mean-field analytical theory for cell shape variability. We find that the AR distribution is described by a single parameter, *α*, containing all the system-specific parameters, leading to a universal relation between *sd* and 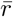. On the other hand, the PDF of *r*_*s*_ is not strictly, but virtually, universal; *k* ≃ 2.5 is a direct consequence of a mathematical property. We also obtain the PDF for the scaled area, *a*, and show that it is not universal, in contrast to what has been proposed elsewhere [36]. We demonstrate that simultaneous measurements of the PDFs for *a* and *r* can reveal the effects of changing physical conditions, such as maturation, on the individual model parameters. We have verified our theory via simulations of two distinct models of a confluent epithelial monolayer: the discrete lattice-based CPM on square and hexagonal lattices and the continuous VM (see Supplementary Material (SM), Sec. S3 for details). Moreover, comparisons with existing experimental data on a wide variety of epithelial systems show excellent agreements. We illustrate that the same parameter describes both statics and dynamics, governs the origin and aspects of universality, and, thus, provides a framework to analyze and compare diverse epithelial systems.

## Analytical theory for the shape variability

Simplified model systems, representing cells as polygons, have been remarkably successful in describing both the static and dynamic aspects of an epithelial monolayer [8, 13, 26, 37–40]. The energy function, ℋ, governing these models is

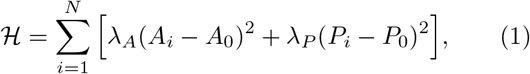

where *N* is the number of cells, the first term constrains area, *A*_*i*_, to a target area *A*_0_, determined by cell height and cell volume, with strength *λ*_*A*_. Experimentally, it has been shown cell height remains almost constant in an epithelial monolayer [13]. The second term describes cortical contractility and adhesion [13, 26, 41]. It constrains the perimeter, *P*_*i*_, to the target perimeter *P*_0_ with strength *λ*_*P*_. The energy function, Eq. (1), can be numerically studied [42] via different confluent models, such as the VM [13, 26], the Voronoi model [27, 37], or more microscopic models such as the CPM [43–45] and the phase-field model [34, 46, 47].

We aim to develop a mean-field theory for a microscopic version, as schematically shown in Fig. 1(a), to calculate the shape variability. The mechanical properties of a cell, for most practical purposes, are governed by a thin layer of cytoplasm just below the cell membrane, the cortex. Therefore, the crucial contribution should come from the perimeter term. We describe the cell perimeter via *n* points, where *l*_*j*_ is the infinitesimal line-element between *j*th and (*j* + 1)th points (Fig. 1a).

**FIG. 1.**
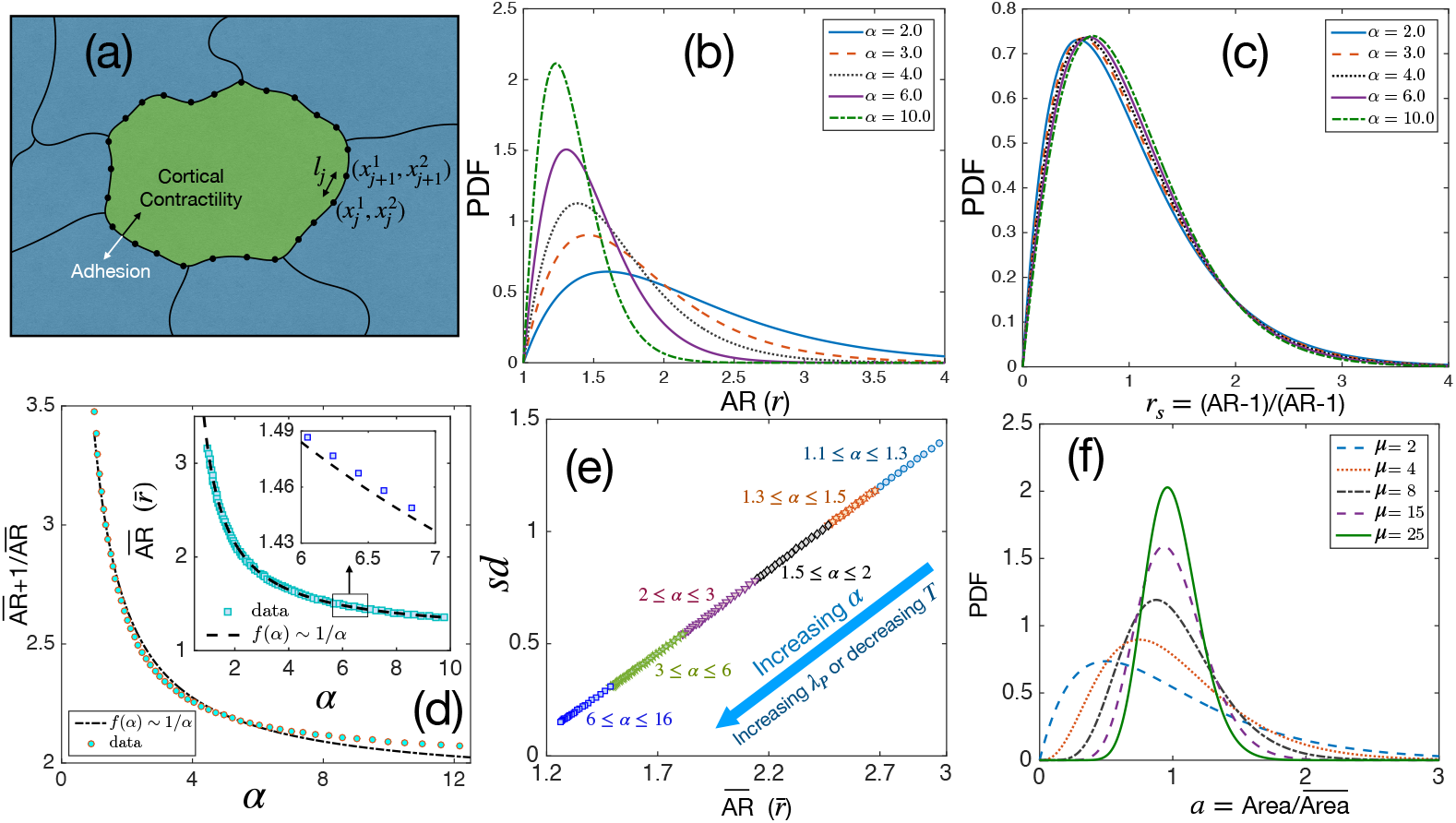
Theoretical results for cell shape variability. (a) Schematic illustration of the mean-field model. Cell perimeter is represented by a set of points (shown for a particular cell); *l*_*j*_ is the distance between two neighboring points. Cortical contractility and adhesion impose competing forces. (b) PDF of aspect ratio, *r*, is governed by the parameter *α*. (c) PDF of the scaled aspect ratio, 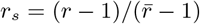 is nearly universal. (d) The PDF of *r*_*s*_ can be universal if 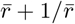 is proportional to 1*/α* (Eq. 7), but there is a slight deviation showing that it is not strictly universal. **Inset:** 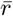 also slightly deviates from the 1*/α* form. Lines are fits with the 1*/α* form. (e) Standard deviation (*sd*) vs 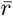 follows a universal relation; the state points move towards the origin as *α* increases. (f) The PDF of *a* follows Gamma distribution with a single parameter, *µ*, Eq. (8).

Now, AR and area can vary independently. We assume that the area constraint is satisfied and ignore the first term in Eq. (1). This assumption is motivated by our simulation results, where the AR distribution is nearly independent of *λ*_*A*_ (Fig. 2f).

**FIG. 2.**
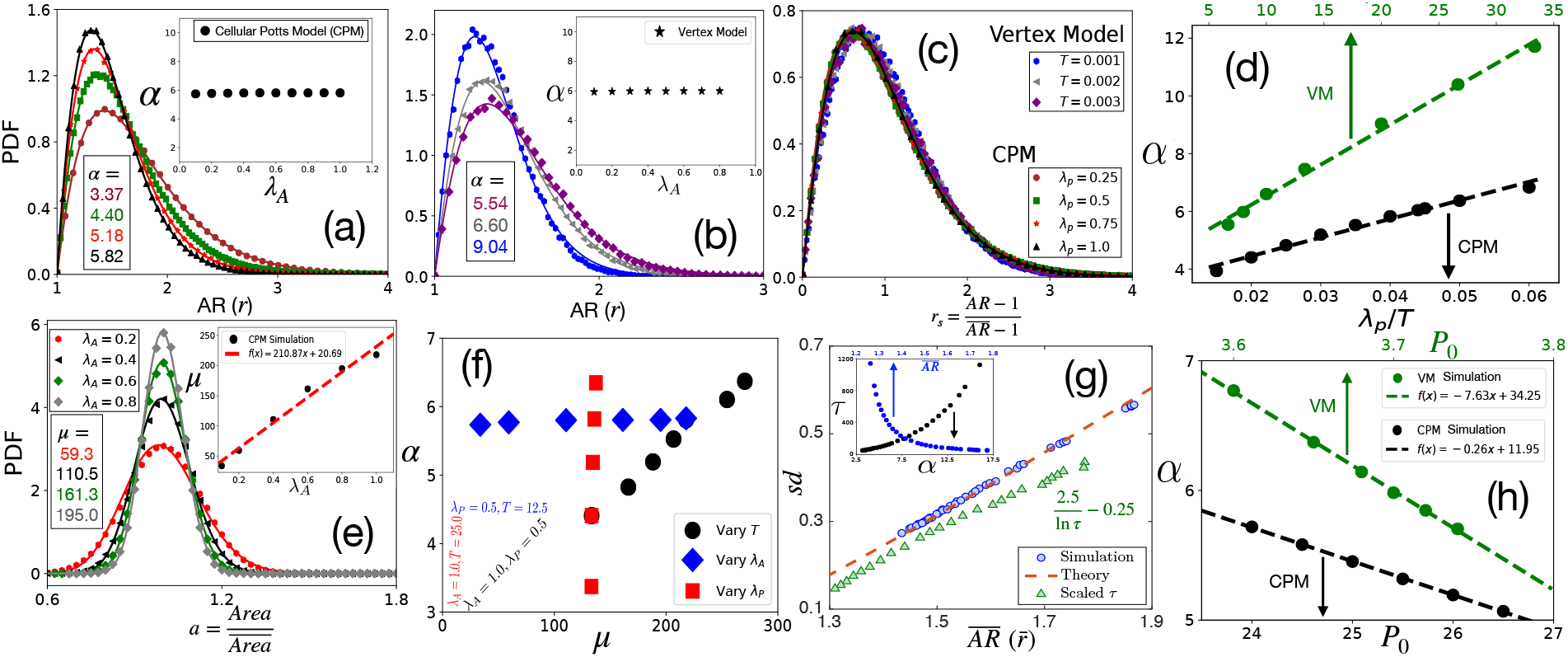
Comparison with simulations. (a) PDF of AR at different *λ*_*P*_ (quoted in c) within the CPM, with *λ*_*A*_ = 1.0, and *T* = 25.0. Symbols are simulation data, and lines represent the fits with Eq. (7). **Inset:** *α* within the CPM at *λ*_*P*_ = 0.5, and *T* = 25.0 with varying *λ*_*A*_ remains almost constant. (b) PDF of *r* within the VM with varying *T* (quoted in c), *λ*_*P*_ = 0.02, *λ*_*A*_ = 0.5, *P*_0_ = 3.7. Lines are fits with Eq. (7) with *α* as shown. **Inset:** *α* as a function of *λ*_*A*_ remains almost constant also within the VM. *T* = 2.5 × 10^−3^ and *λ*_*P*_ = 0.02 and *P*_0_ = 3.7. (c) PDFs for the *r*_*s*_, for the same data as in (a) and (b) for the CPM and the VM, respectively, show a virtually universal behavior. (d) Theory predicts *α* linearly varies with *λ*_*P*_ */T*, simulation data within both the models agree with this prediction. *λ*_*A*_ = 1.0 for the CPM; *P*_0_ = 3.7 and *λ*_*A*_ = 0.5 for the VM simulations. Dotted lines are fits with a linear form. (e) PDF of the scaled area, *a*, within the CPM at different *λ*_*A*_, but fixed *T* = 12.5 and *λ*_*P*_ = 0.5. The lines are fits with Eq. (8) with the values of *µ* as shown. **Inset:** *µ* almost linearly increases with *λ*_*A*_. (f) *α* vs *µ* within the CPM when we have varied only one of the three variables: *T, λ*_*A*_, and *λ*_*P*_. *α* does not depend on *λ*_*A*_, and *µ* does not depend on *λ*_*P*_. (g) Our theory predicts a universal behavior for *sd* vs 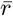, symbols are the CPM data at different parameter values, and the line is our theory (not a fit). We have also plotted the scaled relaxation time, 2.5*/* ln *τ −* 0.25, to show them on the same figure. *τ* increases as 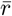 decreases. **Inset:** *τ* as functions of *α* (lower axis) and 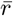 (upper axis). (h) Theory predicts *α* linearly decreases with *P*_0_, this agrees with simulations (symbols); parameters in the CPM: *λ*_*A*_ = 1.0, *λ*_*P*_ = 0.5, *T* = 16.67; the VM: *λ*_*A*_ = 0.5, *λ*_*P*_ = 0.02, *T* = 0.0025. [CPM simulations here are on square lattice, *Ā* = 40, *P*_0_ = 26 in (a-f)].

Next, consider the perimeter term: 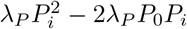; the first term represents contractility, and the second, effective adhesion; we have ignored the constant part. Exact description of adhesion is complex [23, 48, 49]. The VM represents it as a line tension: Σ_⟨ *ij*⟩_Λ_*ij*_ *ℓ*_*ij*_, where *ℓ* _*ij*_ is the length between two consecutive vertices, *i* and *j*, and Λ_*ij*_ gives the line tension. Since the degrees of freedom in the VM are the vertices, considering Λ_*ij*_ constant, we obtain Eq. (1). The constant Λ_*ij*_ implies a regular cell perimeter between vertices. However, the entire cell boundary is in contact with other cells, and the perimeter is irregular in experiments [10, 49]. We take a more general description, where the tension in a line-element *l*_*j*_ (Fig. 1a) is proportional to *l*_*j*_ with strength *P*_0_. Thus, the adhesion part becomes 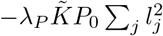, where 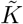 is a constant. On the other hand, the contractile term is proportional to (Σ_*j*_ *l*_*j*_)^2^. To make the calculation tractable, we use the Cauchy-Schwartz inequality [50] and write 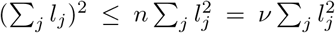, where *ν* ∼ 𝒪 (*n*) is a constant. Then, the perimeter part of ℋ becomes 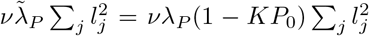, where 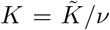. Thus, contractility and adhesion act as two competing effects (Fig. 1a). Since it is unclear how to measure *P*_0_ in experiments, we mainly restrict our discussions on *λ*_*P*_ and *T*. Unless otherwise stated, we assume *P*_0_ remains constant. However, we show in simulations that our mean-field theory captures the variation in *P*_0_ (Fig. 2h). Cell division and apoptosis rates are usually low in epithelial monolayers [8, 12]; for example, they are of the order of 10^−2^ per hour and per day, respectively, for an MDCK monolayer [51]. Nevertheless, we show in the SM (Fig. S4), the general conclusions of the theory remain unchanged even when their rates are significant. We describe the cell perimeter by the vector 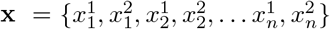, representing a set of *n* points on the perimeter (Fig. 1a). From the preceding discussion, we have 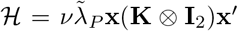, where **x′** is the column vector, the transpose of **x, K**, the Kirchhoff’s matrix [52, 53] with **K**_*ii*_ = 2 and **K**_(*i*−1)*i*_ = **K**_*i*(*i*−1)_ = −1, and **I**_2_, the two-dimensional identity tensor. Thus, we have

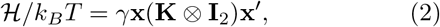

where *γ* = *νλ*_*P*_ (1 − *KP*_0_)*/k*_*B*_*T, k*_*B*_ is the Boltzmann constant, and *T*, the temperature that parameterizes all possible activities, including the equilibrium temperature. To calculate *r*, we first obtain the moment of inertia tensor, a 2 × 2 tensor in spatial dimension two, in a coordinate system whose origin coincides with the center of mass (CoM) of the cell. Diagonalization of this tensor gives the two eigenvalues, Λ_1_ and Λ_2_, that are the squared-radii of gyrations around the respective principal axes. Therefore, 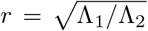, assuming Λ_1_ ≥ Λ_2_. However, a direct calculation of Λ_1_ and Λ_2_ is complex due to their anisotropic natures. It simplifies a bit for the radius of gyration, *s*, around the center of mass, and we have *s*^2^ = Λ_1_ + Λ_2_ [54]. The distribution of *s*^2^ is

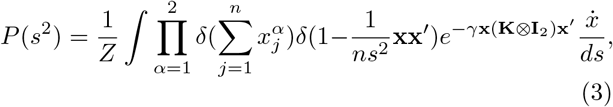

where *Z* is the partition function, 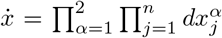, the volume element [52, 53]. Note that the definition of *P* (*s*^2^), Eq. (3), assumes the Boltzmann distribution for perimeter at *T* and accounts for all possible configurations allowed by ℋ. The applicability of an effective *T* at the cellular level might be questionable. But the degrees of freedom here are much more microscopic (Fig. 1a): within the CPM, it’s a pixel, comparable to the resolution of the experimental image processing method. At this level of description, one can safely assume a Boltzmann distribution. Additionally, analyses of the experimental systems [10, 33] suggest the applicability of an effective *T* description. The first *δ*-function in Eq. (3) ensures that the origin coincides with the CoM of the cell; the second *δ*-function selects specific values of *s*^2^, giving the distribution function. We next go to the normal-coordinate system that diagonalizes **K**. Integrating out the coordinates (see SM, Sec. S1), we obtain

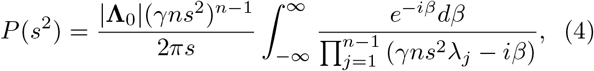

where *λ*_*j*_’s are the non-zero eigenvalues of **K** whose diagonal form, without the zero-eigenvalue, is **Λ**_0_. Taking contour integral of Eq. (4), we obtain

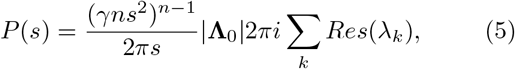

where *λ*_*k*_ are the distinct non-zero eigenvalues of **K** and *Res*(*λ*_*k*_) gives the residue at the pole *λ*_*k*_. Since the cell perimeter must be closed-looped, **K** is a tridiagonal matrix with periodicity. Therefore, the number of zero-eigenvalue must be one, and the lowest degeneracy of the non-zero eigenvalues must be two [52, 55]; as explained below, the value of *k* in *P* (*x, k*) is determined by this second property. The leading-order contribution (SM, Sec. S1) comes from the smallest eigenvalue, *λ*, and is

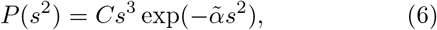

where *C* is a normalization constant and 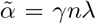. For cells, the two eigenvalues, Λ_1_ and Λ_2_, are not independent since 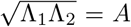, the cell area. Using this relation, we obtain *s*^2^ = *A*(*r* + 1*/r*) and from Eq. (6), we obtain

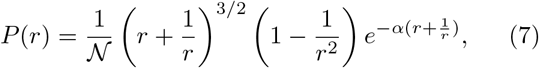

where 𝒩 is the normalization constant: 𝒩= Γ(5*/*2)*/α*^5*/*2^ − *W* (5*/*2)_1_*F*_1_(5*/*2, 7*/*2, *−* 2*α*), where *W* (*x*) = 2^*x*^*/x*, and _1_*F*_1_(*a, b, c*) is the Kummer’s confluent Hypergeometric function [56], *α ∝ λ*_*P*_ (1 − *KP*_0_)*/T*. As detailed in the SM (Sec. S2), Eq. (6), together with the constraint of confluency [57, 58], give the distribution for the scaled area *a* = *A/Ā*, where *Ā* is the average of area, *A*. It is a Gamma distribution, with a single parameter *µ*,

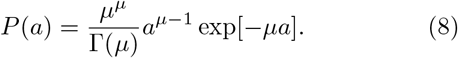

Since *µ* is related to the constraint of confluency, it should be independent of *λ*_*P*_ ; our simulations show that this is indeed true (Fig. 2f). Therefore, *α* and *µ* together can distinguish how the model parameters *λ*_*P*_ and *T* are affected by changing conditions such as maturation. Equation (7) provides a remarkable description, where all the microscopic details of the system enter through a single parameter, *α*; it has profound implications leading to the universal behavior as we now illustrate.

## Aspects of universality

We first show the PDF of the aspect ratio, *P* (*r*), at different values of *α* in Fig. 1(b), *P* (*r*) decays faster as *α* increases, as expected from Eq. (7). The plots look remarkably similar to the experimental results shown in Ref. [10]; we present detailed comparisons with experiments later. Reference [10] has demonstrated that the PDFs of the scaled aspect ratio, 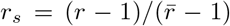, where 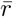 is the ensemble-averaged *r*, across different systems follow a near-universal behavior. We now plot the PDFs of *r*_*s*_ in Fig. 1(c). The PDFs *almost* overlap, but they are not identical. A closer look at Eq. (7) shows that if 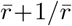 goes as 1*/α*, we can scale *α* out of the equation and obtain a universal scaled distribution. However, as shown in Fig. 1(d), there is a slight deviation in 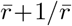 with the functional form of 1*/α*. As shown in the inset of Fig. 1(d), 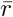 also slightly deviates from the 1*/α* form. This tiny deviation implies that the PDF of *r*_*s*_ is not universal. If one ignores 1*/r* compared to *r*, Eq. (7) becomes a *k*-Gamma function for *r*_*s*_ with *k* = 2.5. However, since *r* ∼ 𝒪 (1), this cannot be a good approximation, and the observed spread of *k* around 2.5 is natural when fitted with this function [10, 33, 34, 59]. On the other hand, since the deviation (Fig. 1d) is minute, the PDFs of *r*_*s*_ for different systems look *virtually* universal. This result is a strong prediction of the theory and, as we show below, is corroborated by available experimental data on diverse epithelial systems.

Although the PDFs of *r*_*s*_ are not strictly universal, there is another aspect, *sd* vs 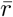, which is universal. We show the 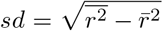 as a function of 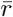 in Fig. 1(e). Since there is only one parameter, *α*, in Eq. (7), it determines both *sd* and 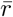. The monotonic dependence of 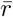 on *α* (inset, Fig. 1(d)) implies a unique relationship between them. Therefore, we can express *α* in terms of 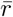 and, in turn, *sd* as a function of 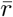. Since there is no other system-dependent parameter in this relation, it must be universal. Note that *α ∝λ*_*P*_ */T* at a constant *P*_0_, thus, *α* increases as *λ*_*P*_ increases or *T* decreases. Both *sd* and 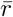 become smaller as *α* increases, and the system on the *sd* vs 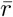 plot moves towards the origin (Fig. 1e). From the perspective of the dynamics, the relaxation time, *τ*, of the system grows as *α* increases [28]. Thus, small 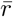 and large *τ*, that is, less elongated cells and slow dynamics, follow each other, and the energy function, Eq. (1), controls both behaviors. Finally, we show some representative PDFs, Eq. (8), at different values of *k* for the scaled area *a* = *A/Ā*. The PDF of *a* has been argued to be universal [36]; our theory shows that although they follow the same distribution, they need not be identical.

## Comparison with simulations

We now compare our analytical theory with simulations of two distinct confluent models: the CPM and the VM. Figures 2(a) and (b) show representative plots for the comparison of the PDFs of *r* within the CPM, and the VM, respectively, where the lines represent fits with Eq. (7). Figure 2(a) shows data with varying *λ*_*P*_, and Fig. 2(b) shows data with changing *T*. As discussed above, our theory predicts almost universal behavior for the PDFs of *r*_*s*_ (Fig. 1c). We plot the simulation data for the PDFs of *r*_*s*_ for both the models at different parameters in Fig. 2(c); the PDFs almost overlap with each other, consistent with the theory (see SM, Fig. S2 for more results). Within our mean-field theory, we have ignored the area term in Eq. (1), arguing that the perimeter term contributes dominantly to the shape. We have obtained *α* at fixed *λ*_*P*_, *P*_0_, and *T* but varying *λ*_*A*_. As shown in the insets of Figs. 2(a) and (b), within both the CPM and the VM, *α* remains almost constant with varying *λ*_*A*_, justifying our assumption. An important prediction of the theory is that the parameter *α*, which governs the behavior of the cell shape variability, is linearly proportional to both *λ*_*P*_ and 1*/T*, hence with *λ*_*P*_ */T*. This prediction also agrees with our simulations (Fig. 2d and SM, Fig. S6). The slopes within the CPM and the VM are different; this possibly comes from the distinctive natures of the two models, but the qualitative behaviors are the same.

We now show the behavior of the PDF of scaled cell area, *a*, in Fig. 2(e) with varying *λ*_*A*_. The lines represent the fits with Eq. (8). Larger values of *λ*_*A*_ ensure the area constraint is more effective and the distributions become sharply peaked around the average area. The inset of Fig. 2(e) shows that *µ* almost linearly increases with *λ*_*A*_. Since the area term is related to the cell height that remains nearly constant and the geometric constraint of confluency, we don’t expect a substantial variation in *λ*_*A*_ in a particular system. However, *µ* also varies with *T* (see SM, Fig. S7). Thus, in contrast to what has been proposed [36], as *T* changes, the PDF of *a*, though well-described by a single parameter *µ* via Eq. (8), can still be different.

Figure 2(f) shows *α* vs *µ* within the CPM when we vary one of the parameters, *λ*_*A*_, *λ*_*P*_, and *T*, keeping the other two fixed. First, when *λ*_*A*_ increases, the value of *µ* increases, but *α* remains almost constant (also see insets of Fig. 2a and b). Next, when *λ*_*P*_ increases, although *α* linearly increases, *µ* remains nearly the same. Finally, both parameters linearly increase with 1*/T* ; since higher *T* implies more fluctuations, decreasing *T* helps both *r* and *a* to become sharply peaked (see SM, Figs. S6, S7 for their specific behaviors, and results within the VM). These results show when *λ*_*A*_ remains constant, varying *λ*_*P*_ and *T* have distinctive effects on *µ* and *α*. These results are significant from at least two aspects: First, *µ* comes from the constraint of confluency (see SM, Sec. S2), which should depend only on the area and be independent of the perimeter. Thus, the *λ*_*P*_ -independence of *µ* validates the phenomenological implementation [57, 58] of this constraint. Second, these results can provide crucial insights regarding the model parameters. Maturation of a monolayer can affect both *λ*_*P*_ and *T*. Additional junctional proteins may be employed during maturation to increase *λ*_*P*_. On the other hand, different forms of activity may reduce decreasing *T*. Since *α* increases linearly with *λ*_*P*_ */T*, AR alone is not enough to determine the dominant mechanism during the maturation process. However, assuming that *λ*_*A*_ remains constant in a particular system, simultaneous measurements of *µ* and *α* allow distinguishing effects of changing physical conditions, such as maturation, on the individual parameters.

We next verify the universal result of the theory: *sd* vs 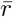. Figure 2(g) shows *sd* vs 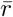 within the CPM; we plot the theoretical prediction by the dotted line for comparison. The theory predicts that the state points move towards the origin as *α* increases (Fig. 1e); this is consistent with our simulations. Since *α ∝ λ*_*P*_ */T*, higher *α* should correspond to slower dynamics. To test this hypothesis, we have simulated the CPM at different *λ*_*P*_, *P*_0_, and *T* to obtain the relaxation time, *τ* (see SM, Sec. S3 for details). From these control parameters, we have calculated *α* and then 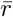 using our theory. We show *τ* as functions of *α* and 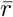 in the inset of Fig. 2(g); it is clear that indeed *τ* grows as *α* increases or 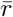 decreases. To show this behavior of *τ* on the same plot as *sd* vs 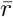, we plot 2.5*/* ln *τ −* 0.25 in the main-figure as a function of 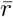. A monolayer fluidizes under compressive or stretching experiments, where cell shape changes, but not cell area [8, 10, 60]. Such perturbations make the cells more elongated, increasing 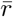; thus, our theory rationalizes the decrease in *τ* associated with fluidization. Finally, we show that our mean-field result that *α* decreases linearly with *P*_0_ agrees with simulations (Fig. 2h). Further, to test our hypothesis that our main results remain unchanged in the presence of cell division and apoptosis, we have simulated the CPM, including these processes. As shown in the SM, Sec. S7, the simulation results justify our hypothesis.

## Comparison with existing experiments

Having shown that our theory agrees well with the simulations, both within the CPM and the VM, we next confront it with the existing experimental data. We first compare the theory with data taken from Ref. [10] for three different confluent cell monolayers: the MDCK cells, the asthmatic HBEC, and the nonasthmatic HBEC. We chose the PDFs at three different times from Figs. 3 (a) and (c) in Ref. [10]. We fit Eq. (7) with the data to obtain *α* and present their values in Table I; the corresponding fits for the MDCK cells are shown in Fig. 3 (a) (see SM, Fig. S8 for the other fits). Table I shows that *α* increases with maturation. Thus, progressive maturation can be interpreted as an increase in either *λ*_*P*_ or 1*/T* or both. The PDFs for *r*_*s*_ corresponding to the MDCK cells are shown in the inset of Fig. 3(a) together with one set of experimental data [10]; note that this is not a fit, yet the theory agrees remarkably well with the data.

**TABLE I.** Values of *α* from fits of Eq. (7) with the PDFs of *r*. The data for the three systems are taken as function of maturation time, in units of h (hours) and d (days). The experimental data are taken from Ref. [10].

| Cell type | Time | Value of $\alpha$ |
| --- | --- | --- |
| MDCK | 32 h | 7.535 |
|  | 47 h | 10.998 |
|  | 63 h | 12.496 |
| Asthmatic HBEC | 6 d | 3.060 |
|  | 14 d | 4.472 |
|  | 20 d | 5.389 |
| Non-asthmatic HBEC | 6 d | 4.798 |
|  | 14 d | 7.358 |
|  | 20 d | 9.602 |

**FIG. 3.**
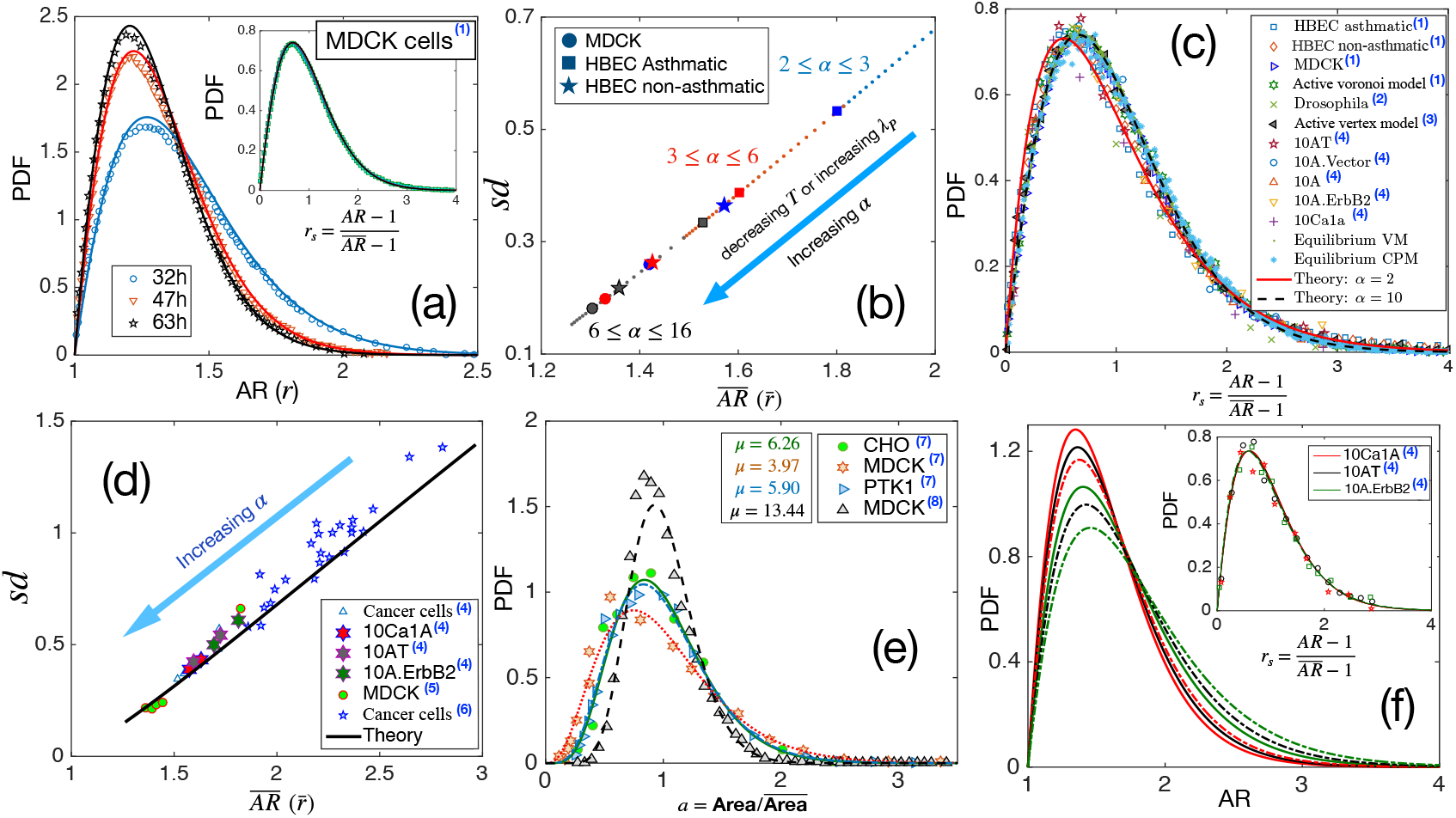
Comparison with existing experiments. (a) PDF for AR. Symbols are experimental data at three different times for MDCK cells, and lines represent the fits with our theory, Eq. (7). The values of *α* are quoted in Table I. **Inset:** PDF of the *r*_*s*_, the lines are theory, and the symbols are data. (b) Using the values of *α*, obtained for the three sets of data as quoted in Table I, we obtain *sd* vs 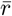 using our theory. Different symbols represent the types of the systems, and the colors blue, red, and black represent early, intermediate, and later time data, respectively. With maturation, the system moves towards lower 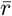 and smaller *sd*. (c) PDF for *r*_*s*_ for a wide variety of systems, shown in the figure, seems to be virtually universal, consistent with our theory. (d) The theory predicts a universal relation for *sd* vs 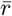. Symbols are data for different systems, and the line is our theoretical prediction. (e) PDF of the scaled area for different epithelial systems and the lines represent the fits with Eq. (8). (f) Predictive power of the theory: we use the 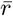 for the three sets of cells from Ref. [33], as marked by the hexagons in (d), and obtain the corresponding values of *α* and obtain the PDFs for *r* using Eq. (7). The colors correspond to the type of cells in (d), and the continuous line corresponds to the lower 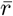 data. **Inset:** Lines are the theoretical PDFs for *r*_*s*_ using the values of *α* for different cells, and the symbols show the experimental data. We have collected the experimental and some of the simulation data from different papers. Data taken from other papers are marked with a blue superscript in the legends. The sources are as follows: (1) [10], (2) [32], (3) [31], (4) [33], (5) [61], (6) [35], (7) [36] and (8) [51].

We next calculate *sd* as a function of 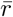 using the values of *α*, noted in Table I for the three systems. They are shown in Fig. 3(b) along with the theory prediction. With maturation, the state points move towards lower 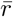, represented by the arrow in Fig. 3(b). As shown in Fig. 2(g), larger *α* corresponds to a system with higher *τ*. Thus, with maturation, as the PDFs become sharply peaked, as the cells look more roundish and 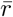 becomes smaller, the system becomes more sluggish. This effect of maturation is the same in all the systems (Fig. 3b) and agrees with the interpretation presented in Ref. [10]. We have also examined that the theoretical prediction of *sd* vs 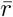 agrees well with the experimentally measured values shown in Ref. [10].

Our theory predicts that the PDF for *r*_*s*_, though not strictly universal, should be almost the same for different systems (Fig. 1c). This prediction is a consequence of a crucial aspect of the theory: all the system-specific details enter via a single parameter, *α* in Eq. (7). As shown in Fig. 1(d), 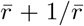 deviates slightly from the behavior 1*/α*. This slight deviation implies that the PDF for *r*_*s*_ can not be strictly universal and manifests as a variation in *k* when the PDF is fitted with the *k*-Gamma function in different experiments and simulations [10, 33, 34, 59, 62]. Nevertheless, since the deviation in Fig. 1(d) is very weak, the values of *k* are very close to each other. Therefore, the PDFs for *r*_*s*_ for diverse epithelial systems–in experiments, simulations, and theory– should be virtually universal. To test this prediction, we have collected existing experimental and simulation data on different systems and show the PDFs of *r*_*s*_ on the same plot in Fig. 3(c). The variety in our chosen set is spectacular: it consists of different cancer cell lines [33], both asthmatic and non-asthmatic HBEC cells, MDCK cells [10], Drosophila wing disk [62], simulations data on both active [59] and equilibrium versions of the VM, the active Voronoi model [10], and the CPM. Yet, the PDFs, as shown in Fig. 3(c), look virtually universal and in agreement with our analytical theory.

Additionally, our theory predicts a strictly universal *sd* vs 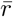. Since this relation does not have any system-specific details, data across diverse confluent monolayers must follow a universal relationship. We have collected existing experimental data for several systems: cancerous cell lines [33], human breast cancer cells [35], and a jammed epithelial monolayer of MDCK cells [61]. Figure 3(d) shows the experimental data together with our theoretical prediction; the agreement with our theory, along with the aspect of universality, is truly remarkable. As *α* increases, dynamics slows down, and the points on this plot move towards lower 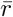. This result is consistent with the finding that cell shapes are more elongated and variable as the dynamics become faster in different epithelial systems [8, 10].

We have argued that simultaneous measurements of the PDFs of cell area and AR distinguish the effects of maturation on the two key parameters: *λ*_*P*_ and *T*. The argument relies on the negligible effect of the perimeter constraint on *µ* (Eq. 8). We now show a comparison of our theoretical result for the PDF of *a* with existing experiments. Figure 3(e) shows experimental data for four different systems [36, 51] and the corresponding fits of Eq. (8). Unlike what has been proposed, that epithelial monolayers have a universal area distribution [36], we find, in agreement with experiments, the distribution can vary, although the functional form remains the same.

What are the implications of these universal aspects of cell shape variability and our theory? Cell shape controls several crucial biological functions such as the mitotic-orientation [3, 15, 16] and cell fate [17, 18, 20]. Our theory shows that the microscopic system properties are encoded via a single parameter, *α*. Consequently, knowledge of one of the observables, such as 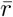, contains the information of the entire statistical properties in a monolayer. We now illustrate this predictive aspect of the theory. Experimental measurement of an average property is usually less complex and more reliable. We have collected the data for 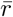 from the supplementary material of Ref. [33] for three different systems: 10Ca1A, 10AT, and 10A.ErbB2, shown by the hexagons in Fig. 3(d). From these average values, we obtain *α*, which we use to theoretically calculate the PDFs for *r*, as shown in Fig. 3(f). The inset of Fig. 3(f) shows our theoretical PDFs for *r*_*s*_, together with the corresponding experimental data for comparison. The excellent agreement demonstrates that cell shape variability results from the geometric constraint imposed by the energy function, Eq. (1), and is not a choice but inevitable for such systems. This result, we believe, will foster analysis of diverse epithelial systems to understand the interrelation between geometric properties and biological functions within a unified framework.

## Discussion

We have obtained a mean-field theory for cell shape variability through the energy function ℋ, Eq. (1) [13, 26–28, 37]. The geometric restriction of confluency is a strong constraint on cell area. Considering that it is satisfied and that the cell cortex, described by the perimeter term, is crucial in determining the cell shape, allowed us to ignore the area term and obtain the distribution for AR. Recent experiments and simulations have shown that cell shape variability is virtually universal in a confluent epithelial monolayer [10, 33–35]; our work provides the theoretical basis for such behavior. Contrary to common beliefs [10, 33], the jamming transition is not related to the cell shape distribution. We have analytically obtained the functional form for cell shape variability and find that the microscopic system properties enter the distribution via a single parameter, *α*: this leads to the universal behavior for *sd* vs 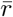 and a virtually universal distribution for the scaled aspect ratio *r*_*s*_. Our theoretical results are verified in simulations within the CPM, on the square and hexagonal lattices, and the VM. They also agree remarkably well with the existing experiments. Thus far, the PDF of *r*_*s*_ has been fitted with a *k*-Gamma function with *k* being around 2.5. We show that the slight variation in *k* comes from the fact that the PDF is not strictly but almost universal. On the other hand, *k* ≃ 2.5 is a direct consequence of a mathematical property: the lowest degeneracy of the eigenvalues of the connectivity matrix being two for a closed-looped object, here the perimeter.

A better understanding of the connection between the theoretical parameters and different system properties is crucial to exploit the universal aspects for deeper insights. Since all of the parameters combine into *α*, the effects of changing physical conditions on the individual parameters are difficult to determine from the measurements of AR alone. Within our theory, *λ*_*P*_ describes the cortical properties, and *T* parameterizes different biological activities, including temperature. These are effective parameters, and their direct measurement in biological systems is impractical. Our theory provides an indirect way to estimate these parameters. The cell area in a confluent system is geometrically constrained. We have used the phenomenological implementation of the constraint of confluency and obtained the PDF for *a* [57]. It is a Gamma function, described by a single parameter *µ*. Reference [36] has proposed that the PDF of *a* is universal in various epithelial monolayers. However, we show that though the area distribution follows the same function, there is a variation in *µ*. Our work connects *µ* to the microscopic model parameters of Eq. 1. In particular, *µ* should be independent of *λ*_*P*_, whereas *α* varies linearly with both *λ*_*P*_ and 1*/T*. This distinction, assuming *λ*_*A*_ remains constant, allows inferring the effects of maturation on the individual model parameters. We have neglected cell division, growth, and apoptosis in our theory. However, the consequences of these processes, and the effects of chemical gradients [63], should also enter the distribution via *α*, as the algebraic part comes from the topological property that remains the same. We have included them in our simulations, and the results support this expectation (SM, Sec. S7); this implies that the main predictions of the theory remain valid even in the presence of these processes.

Our work demonstrates that a single parameter describes both the cell shape statistics and the dynamics. We have shown in our simulations that the relaxation time, *τ*, grows as *α* increases or 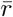 decreases. Experiments on confluent cellular monolayer have also reported similar results [8, 10, 33]. Thus, as cells become more compact and their shape variability reduces, the monolayer becomes more sluggish. This result may have far-reaching consequences. Most experiments usually measure 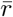 and analyze biological functions via 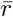 [3, 15]. Our theory implies that such knowledge contains a wealth of information. One can obtain *α* from 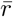, and all other properties, such as the distribution, the standard deviation, and the dynamics, can be analytically calculated.

Whether this correlation between the statics and dynamics persists in all regimes, including the glassy-regime, remains an open question. Additionally, in biology, it is well-recognized that different levels of organizations are mechanistically related. One fundamental open question is how molecular-level events are related to cellular machines that control the cell shape [64]. The striking predictability, demonstrated by our theory, where 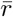 determines the PDF, shows that the statistical distribution of cell shape is unavoidable. How do different cells respond to this inevitable distribution? Is cellular response similar across diverse systems? How is it related to organ-level morphogenesis? Having a single parameter that describes the static and dynamic aspects at the cellular level should help to compare and analyze different systems and answer these questions.

In conclusion, we have developed a mean-field theory, considering cortical contractility and adhesion are crucial in determining the shape. We have analytically derived the cell shape variability, characterized via the AR. The PDF for AR is described by a single parameter, *α*. As a result, *sd* vs 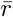 becomes universal, and the PDF for the scaled aspect ratio, *r*_*s*_, is virtually universal. The analytical form for the PDF of *r*_*s*_ can be roughly approximated as a *k*-Gamma distribution [30] that has been fitted with the existing experimental data [10, 33]; however, the distribution is unrelated to the jamming transition, in contrast to the general belief. The near-universal value of *k* ∼ 2.5 is the consequence of a mathematical property, and the variation results from the fact that the PDFs are not strictly universal. *α* can also provide information on dynamics. In addition, simultaneous measurements of the PDFs of *a* and *r* can relate the model parameters with the changing physical properties. Such a relationship should further strengthen the link between the energy function, Eq. (1), and an epithelial monolayer. Having a single parameter for the statistical and dynamical aspects of an epithelial monolayer should foster a detailed comparison of diverse systems, help to analyze the relation of biological functions to shapes, and illuminate the mechanisms of cellular response to the inevitable shape variability.

## Acknowledgements

We thank Mustansir Barma and Sumedha for many enlightening discussions and Sam. A Safran for comments. We acknowledge support of the Department of Atomic Energy, Government of India, under Project Identification No. RTI 4007.

## Notes

### Competing Interest Statement

The authors have declared no competing interest.

