## Supplementary Material for "The origin of universal cell shape variability in a confluent epithelial monolayer"

In this supplementary material, we first provide the details of the analytical derivation of the probability distribution functions (PDFs) of the aspect ratio (AR) in Sec. S1, and the scaled area, $a$, in Sec. S2. We next provide the details of the simulations in Sec. S3, and explain the procedure to calculate the AR in our simulations in Sec. S4. We then present data of the CPM on the hexagonal lattice in Sec. S5, and show additional data in Sec. S6, supporting the claim that AR does not depend on $\lambda_A$. Section S7 shows simulation results when we include cell division and apoptosis. We present the PDF of $a$ within the vertex model in Sec. S8, show the dependence of the parameter $\alpha$ on $\lambda_P$ and $T$ in Sec. S9. The $T$-dependence of $\mu$ is shown in Sec. S10 and, finally, the fits with the experimental data of HBEC cells are shown in Sec. S11.

### S1. Details of the derivation

As shown in the main text, defining $\mathbf{x} = \{x_1^1, x_1^2, x_2^1, x_2^2, \dots x_n^1, x_n^2\}$ as a particular configuration of the perimeter of a specific cell, we can write the energy in units of $k_BT$ for this cell as

$$\mathcal{H}/k_BT = \gamma \mathbf{x}(\mathbf{K} \otimes \mathbf{I}_2)\mathbf{x}' \tag{S1}$$

where $k_BT$ is Boltzmann constant times temperature, $\gamma = \nu\lambda_P(1 - KP_0)/k_BT$ , $\mathbf{K}$ is the $n$-dimensional Kirchoff's matrix with $\mathbf{K}_{ii} = 2$ and $\mathbf{K}_{(i-1)i} = \mathbf{K}_{i(i-1)} = -1$. $\mathbf{I}_2$ is the two-dimensional identity tensor and $\otimes$ denotes the tensor product. $\mathbf{x}'$ is a column vector, the transpose of $\mathbf{x}$. Then, the distribution of the radius of gyration [1, 2] can be written as

$$P(s^2) = \frac{1}{Z}\int \prod_{\alpha=1}^{2} \delta(\sum_{j=1}^{n} x_j^\alpha)\delta(1 - \frac{1}{ns^2}\mathbf{x}\mathbf{x}') \exp(-\gamma \mathbf{x}(\mathbf{K}\otimes\mathbf{I}_2)\mathbf{x}')\frac{\dot{x}}{ds} \tag{S2}$$

where the volume element $\dot{x}$ is defined as $\dot{x} = \prod_{\alpha=1}^{2}\prod_{j=1}^{n} dx_j^\alpha$. The first $\delta$-function in Eq. (S2) coincides center of mass with the origin of the coordinate system. The squared radius of gyration is $s^2 = n^{-1}\mathbf{x}\mathbf{x}'$ and the second $\delta$-function in Eq. (S2) ensures that the correct values of $s$ are chosen for the distribution. $Z$ is the partition function of the system. Since the radius of gyration does not depend on the coordinate system, we are allowed to chose one that diagonalizes $\mathbf{K}$. Say the diagonal matrix is $\mathbf{\Lambda}$, and $\mathbf{q}$ represents the normal coordinates in this system.

The radius of gyration can be defined as the root-mean-square distance of different parts of a system either from its center of mass or around a given axis. We have designated the former as $s$, defined as

$$s = \sqrt{\frac{1}{N}\sum_{i=1}^{N}(\mathbf{x}_i - \mathbf{x}_{CM})^2}, \tag{S3}$$

where $N$ is the total volume element in the system with coordinates $\mathbf{x}_i$, and $\mathbf{x}_{CM}$ is the center of mass (CoM) of the system. The other two radii of gyration can be defined around the two principal axes (since we are in spatial dimension two) passing through the CoM. We calculate these two radii of gyration by writing the inertia tensor in a coordinate system whose origin coincides with the CoM and diagonalizing the tensor. The eigenvalues, $\Lambda_1$ and $\Lambda_2$, give the square of the radii of gyration. Thus, the aspect ratio, $r$, is obtained as $r = \sqrt{\Lambda_1}/\sqrt{\Lambda_2}$. Due to the anisotropic nature of $\Lambda_1$ and $\Lambda_2$, a direct calculation for their distributions is more complex than that of $s$. We first calculate the distribution for $s$ and then, using this result, obtain the distribution of $r$.

Equation (S2) can be written in the normal coordinate system as

$$P(s^2) = \frac{1}{Z}\int \prod_{\alpha=1}^{2} \delta(q_n^\alpha)\delta(1 - \frac{1}{ns^2}\mathbf{q}\mathbf{q}') \exp(-\gamma \mathbf{q}(\mathbf{\Lambda} \otimes \mathbf{I}_2)\mathbf{q}')\frac{\dot{q}}{ds} \tag{S4}$$

where we have used $q_n^\alpha \propto \sum x_j^\alpha$ that corresponds to the zero-eigenvalue mode of the matrix. Integrating over $q_n^\alpha$, we get rid of this zero-eigenvalue that gives translation. Thus,

$$P(s^2) = \frac{1}{Z}\int \delta(1 - \frac{1}{ns^2}\mathbf{q}_0\mathbf{q}_0') \exp(-\gamma \mathbf{q}_0(\mathbf{\Lambda}_0 \otimes \mathbf{I})\mathbf{q}_0')\frac{\dot{q}_0}{ds} \tag{S5}$$

where we have defined $\mathbf{q}_0$ as the $2(n-1)$ dimensional vector excluding the coordinates corresponding to the zero-eigenvalue. The normalization factor, $Z$, can be calculated exactly through the integration as

$$Z \equiv \int \exp(-\gamma \mathbf{q}_0(\mathbf{\Lambda}_0 \otimes \mathbf{I}_2)\mathbf{q}_0')\dot{q}_0 = \left(\frac{\pi}{\gamma}\right)^{(n-1)} |\mathbf{\Lambda}_0|^{-1}. \tag{S6}$$

Note that the integration in the calculation of $P(s^2)$ is around the boundary of the cell; to separate out the radial part, we now write the volume element in polar coordinate $\mathbf{u}$: $\mathbf{q}_0 = n^{1/2}s\mathbf{u}$. Then $\dot{q}_0 = n^{(n-1)}s^{2(n-1)-1}ds\dot{u}$ and $\mathbf{q}_0\mathbf{q}_0'/ns^2 = \mathbf{u}\mathbf{u}' = 1$. Thus, we obtain from Eq. (S5)

$$P(s^2) = \mathcal{A}\int_{-\infty}^{\infty} d\beta \int e^{-i\beta} e^{-\gamma ns^2 \mathbf{u}[(\mathbf{\Lambda}_0 - \frac{i\beta}{n\gamma s^2}\mathbf{I}_{n-1})\otimes \mathbf{I}_2]\mathbf{u}'}\dot{u}, \tag{S7}$$

with $\mathcal{A} = \left(\frac{\gamma}{\pi}\right)^{(n-1)} |\mathbf{\Lambda}_0| \frac{1}{2\pi} n^{(n-1)} s^{2(n-1)-1}$. $\mathbf{I}_{n-1}$ is the identity matrix of rank $n-1$. Carrying out the integration over $\mathbf{u}$, we obtain

$$P(s^2) = \mathcal{A}\int_{-\infty}^{\infty} d\beta e^{-i\beta} \frac{\left(\frac{\pi}{\gamma ns^2}\right)^{(n-1)}}{\left|\mathbf{\Lambda}_0 - \frac{i\beta}{\gamma ns^2}\mathbf{I}\right|}. \tag{S8}$$

Using the value of $\mathcal{A}$, we obtain

$$P(s^2) = \frac{1}{2\pi s}\int_{-\infty}^{\infty} d\beta \frac{e^{-i\beta}}{\prod_{j=1}^{n-1}\left(1-\frac{i\beta}{\gamma n s^2\lambda_j}\right)}$$
$$= \frac{(\gamma n s^2)^{n-1}}{2\pi s}|\mathbf{\Lambda}_0|\int_{-\infty}^{\infty} d\beta \frac{e^{-i\beta}}{\prod_{j=1}^{n-1}(\gamma n s^2\lambda_j - i\beta)}, \tag{S9}$$

where $\lambda_j$'s are the eigenvalues of $\mathbf{K}$. The integral in Eq. (S9) can be performed via the contour integral and the resultant solution can be written as

$$P(s^2) = \frac{(\gamma n s^2)^{n-1}}{2\pi s}|\mathbf{\Lambda}_0| 2\pi i \sum_k Res(\lambda_k), \tag{S10}$$

where $\lambda_k$ are the distinct eigenvalues of $\mathbf{K}$ and $Res(\lambda_k)$ gives the residue at the pole $\lambda_k$. As we show below, the residues will have a term $\exp[-n\gamma s^2\lambda_k]$ and in the limit $s^2 \to \infty$, only the smallest $\lambda_k$ will contribute.

Since the cell perimeter must be closed-looped, $\mathbf{K}$ is a tridiagonal matrix with periodicity. Therefore, the number of zero-eigenvalue must be one, and the lowest degeneracy of the non-zero eigenvalues must be two [1–4]. We have already integrated out the coordinate corresponding to the zero-eigenvalue. Let us designate the lowest non-zero eigenvalue as $\lambda$. The pole corresponding to $\lambda$ is located at $\beta = -i\gamma n s^2\lambda$, and of order 2. Thus, we obtain the residue as

$$\text{Res} = \frac{d}{d\beta}\left[\frac{e^{-i\beta}}{\prod_{j=1}^{n-3}(\gamma n s^2\lambda_j - i\beta)}\right]_{\beta=-i\gamma n s^2\lambda}. \tag{S11}$$

Let's first take the derivative, with respect to $\beta$, of the numerator and write part of the residue as

$$\text{term1} = -i\frac{e^{-\gamma n s^2\lambda}}{(\gamma n s^2)^{n-3}\prod_{j=1}^{n-3}(\lambda_j - \lambda)}. \tag{S12}$$

Next, differentiating the denominator, we obtain the other part of the residue as

$$\text{term2} = e^{-i\beta}\left[\frac{i}{(\gamma n s^2\lambda_1 - i\beta)^2\prod_{j=2}^{n-3}(\gamma n s^2\lambda_j - i\beta)}\right.$$
$$\left.+\frac{i}{(\gamma n s^2\lambda_2 - i\beta)^2\prod_{j=1,j\neq 2}^{n-3}(\gamma n s^2\lambda_j - i\beta)} + \ldots\right]\Bigg|_{\beta=-i\gamma n s^2\lambda}$$
$$= i\frac{e^{-\gamma n s^2\lambda}}{(\gamma n s^2)^{n-2}\prod_{j=1}^{n-3}(\lambda_j - \lambda)}\left[\frac{1}{\lambda_1-\lambda}+\frac{1}{\lambda_2-\lambda}+\frac{1}{\lambda_3-\lambda}+\ldots\right]. \tag{S13}$$

A comparison of term1 and term2, given by Eqs. (S12) and (S13), respectively, shows that there is an extra factor of $s^2$ in the denominator of term2. Thus, term2 can be ignored compared to term1. Therefore, we obtain the distribution function for $s^2$ as

$$P(s^2) = \frac{|\mathbf{\Lambda}_0| n^2\gamma^2}{\prod_{j=1}^{n-3}(\lambda_j - \lambda)} s^3 e^{-\gamma n\lambda s^2} \equiv C s^3 e^{-\tilde{\alpha} s^2} \tag{S14}$$

where $C$ is the normalization constant that we will fix later, and $\tilde{\alpha} = \gamma n \lambda$. Note that the lowest eigenvalue for the $n$-dimensional Kirchoff's matrix is proportional to $1/n$, thus $n\lambda \sim \mathcal{O}(1)$.

Now, $s^2 = \Lambda_1 + \Lambda_2$ and the aspect ratio $r = \sqrt{\Lambda_1}/\sqrt{\Lambda_2}$. Moreover, we have $\sqrt{\Lambda_1}\sqrt{\Lambda_2} = A$, where $A$ is the average area. Since the distribution of cell area is sharply peaked (Fig. 2e in the main text), as the cell division and apoptosis are slow processes, $A$ can be taken as a constant. Therefore, using the last two relations in the first, we obtain $s^2 = A(r + 1/r)$. Thus, we obtain from Eq. (S14), the distribution of $r$ as

$$P(r) = \frac{1}{\mathcal{N}} \left( r + \frac{1}{r} \right)^{3/2} (1 - \frac{1}{r^2}) e^{-\alpha(r + \frac{1}{r})} \tag{S15}$$

where the condition that total probability must be unity sets the normalization constant $\mathcal{N}$ as

$$\mathcal{N} \simeq \frac{\Gamma(5/2)}{\alpha^{5/2}} - W(5/2)\ {}_1F_1(5/2, 7/2, -2\alpha), \tag{S16}$$

where, $W(x) = 2^x/x$ and ${}_1F_1(a, b, c)$ is the Kummer's confluent Hypergeometric function [5]. Using the value of $\gamma$, we obtain $\alpha \propto \lambda_P(1 - KP_0)/T$. For the analysis of the theory, a symbolic software, such as "Mathematica" [6] is helpful. We note here that the relation between $sd$ and $\bar{r}$ is quite complex, though for most practical purposes it can be taken as a straight line: $sd \simeq 0.71\,\bar{r} - 0.75$.

### S2. Distribution for area

Equation (S14) gives the distribution of the radius of gyration of different cells in a monolayer. Using this equation, we now derive the distribution of area, $A$. Since, AR and $A$ can vary independently, we can use $s^2 \propto A$, as we are interested in the functional form. Therefore, Eq. (S14) gives

$$P(A) \sim A^{3/2} \exp[-\beta A] \tag{S17}$$

where $\beta$ is a constant related to $\tilde{\alpha}$. Note that we ignored the area term in the energy function, $\mathcal{H}$, in the derivation of Eq. (S14). In a confluent cellular monolayer, the individual cell area must obey the strong geometric constraint of confluency; thus, it becomes crucial for the area distribution. However, mathematically imposing this constraint is a challenging geometric problem for random patterns, and no exact result exists yet. Weaire *et al* [7] proposed a phenomenological implementation of this constraint as a polynomial in the area; this remains one of the simplest possible ways to date to deal with this constraint [8]. Keeping only one term of this polynomial for simplicity, we can write this constraint as $f(A) \sim A^{\nu}$ [7, 8]. Using this in Eq. (S17), we obtain

$$P(A) \sim A^{\mu - 1} \exp[-\beta A], \tag{S18}$$

where we have defined $\mu = \nu + 5/2$. From Eq. (S18), we obtain the average area, $\bar{A} = \mu/\beta$. Thus, we obtain the normalized distribution for the scaled area, $a = A/\bar{A}$, as

$$P(a) = \frac{\mu^\mu}{\Gamma(\mu)} a^{\mu-1} \exp[-\mu a], \tag{S19}$$

this is the well-known $k$-Gamma function, defined in Ref. [9], and usually denoted with the variable $k$. This same function has been used in fitting the scaled AR, $r_s$, data in different existing experiments and simulations. Therefore, to avoid confusion with $k$, which is obtained fitting the $r_s$ data, we have used $\mu$ to define the distribution function for $a$. Since $\mu$ comes from the constraint of confluency, it should be independent of $\lambda_P$. Our simulation results within both the CPM and the VM (Fig. 2(f), in the main text) support this hypothesis.

### S3. Simulation details

We have verified our theory via simulations of two distinct models: the CPM and the VM. In both the simulations, we have used the energy function $\mathcal{H}$, given in Eq. (1) in the main text. The CPM is a lattice-based model. The underlying lattice has some effects on the quantitative aspects of the model; for example, on the square lattice, the object with the minimum possible perimeter for a given area is a square. However, the qualitative behaviors are independent of the lattice. To ascertain that the simulation results are independent of the lattice, we have simulated the CPM on two different lattices: the square lattice and the hexagonal lattice.

For the glassy dynamics, the geometric restriction leads to two different regimes, the low-$P_0$, and the large-$P_0$ regime. However, the large-$P_0$ regime characterizes quite large adhesion compared to the cortical contractility; this regime, we feel, is not relevant for the experiments. Therefore, we present most results when $P_0$ is not too large. Here, we briefly discuss the different models.

**Cellular Potts Model (CPM):** In CPM, dynamics proceeds by stochastically updating one boundary lattice site at a time. In a Monte-Carlo simulation, we accept a move with a probability, $P(\sigma \to \sigma') = e^{-\frac{\Delta\mathcal{H}}{T}}$, where we have set Boltzmann constant $k_B$ to unity, $\Delta\mathcal{H}$ is the change in energy going from one configuration to the other. $\sigma$ and $\sigma'$ are the cell indices of the current cell and the target cell indices, respectively. Unit of time refers to $M$ such elementary moves, where $M$ is the total number of lattice sites in the simulation.

*With square lattice* : We have implemented CPM with square lattice in Fortran 90 and followed a *Connectivity Algorithm* developed in [10] to prevent fragmentation of cells [11]. We have chosen a simulation box of size $120 \times 120$ with 360 total cells in the system. The average area of cells in the system is 40, and the minimum possible perimeter on a square lattice with this area is 26. Unless otherwise specified, we always start with an initial configuration where each cell is rectangular with a size $5 \times 8$. We first equilibrate the system for $8 \times 10^5$ MC time steps before collecting the data of aspect ratio. The variables that characterize the system are $\lambda_A$, $\lambda_P$, $P_0$, and $T$. We have ignored cell division and apoptosis, as discussed in the main text. However, to test the effects of these processes, we have included them within this model and present a specific set of results in Sec. S7.

To obtain the relaxation time, $\tau$, we have calculated the overlap function, $Q(t)$, defined as

$$Q(t) = \overline{\frac{1}{N}\sum_{\sigma=1}^{N}\langle W(a - |\mathbf{X}_{cm}^{\sigma}(t+t_0) - \mathbf{X}_{cm}^{\sigma}(t_0)|)\rangle_{t_0}}, \tag{S20}$$

where $X_{cm}^{\sigma}(t)$ is the center of mass at time $t$ of a cell with index $\sigma$, $W(x)$ is a Heaviside step function

$$W(x) = \begin{cases} 1 & \text{if } x \geq 0 \\ 0 & \text{if } x < 0, \end{cases} \tag{S21}$$

$\langle \ldots \rangle_{t_0}$ denotes averaging over initial times and the overline implies ensemble average. The parameter $a$ is related to the vibrational motion of the particles (here cells) inside the cage formed by its neighbors. We have set $a = 1.12$ [11]. We have taken 50 $t_0$ averaging and 20 configurations for ensemble averaging.

*With Hexagonal lattice* : To simulate CPM on the hexagonal lattice, we have used an open-source application CompuCell3D [12, 13]. CPM is the core of CompuCell3D. We have simulated the 2D confluent cell monolayer with periodic boundary conditions using the same energy function, Eq. (1), in the main text. In this simulation, we have taken $128 \times 128$ simulation box size with 361 cells. Initially, Cells are of three different types of areas and perimeters. We have started with a rectangular slab of the cell monolayer. We first equilibrate the system for $10^5$ MC time steps before collecting the data. Fragmentation is allowed in these simulations; however, we have restricted ourselves in the low $T$ regime, where fewer cells are fragmented. While calculating the PDFs, we have eliminated the fragmented cells.

**Vertex Model (VM):** Vertex models(VM) can be viewed as the continuum version of the CPM. Within VM, vertices of the polygons are the degrees of freedom, and cell edges are defined as straight lines (or lines with a constant curvature) connecting between vertices [14, 15]. Within the Monte-Carlo simulation [16], dynamics proceeds by stochastically updating each vertex position of all the cells by a small amount $\delta r$. We accept the move with a probability $P(\mathcal{C} \to \mathcal{C}') = e^{-\frac{\Delta\mathcal{H}}{T}}$ where we have set Boltzmann constant $k_B$ to unity, $\Delta\mathcal{H}$ is the change in energy going from $\mathcal{C} \to \mathcal{C}'$.

Initially, we start will 1024 cells having equal area (= 1.0) and perimeter (= 3.72). We have equilibrated the system for $2 \times 10^6$ MC steps before the start of collecting data.

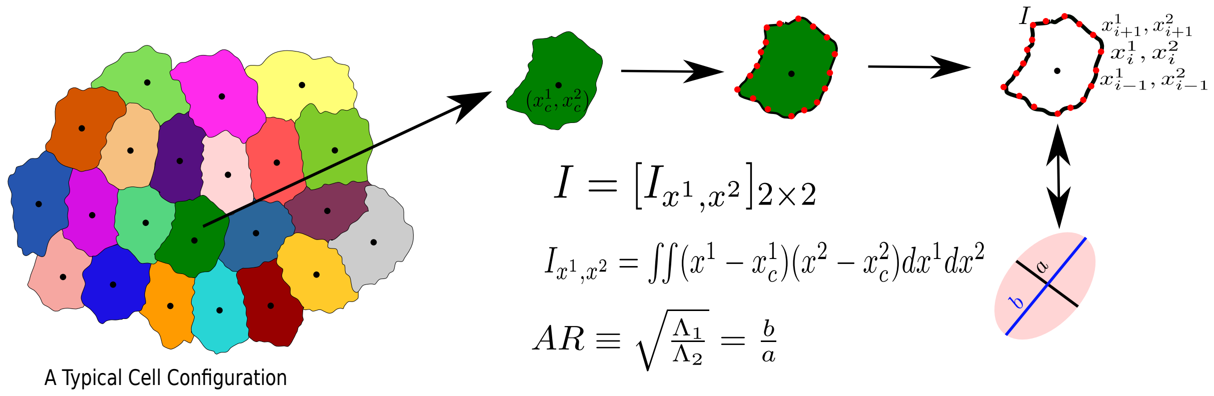


FIG. S1. The process to calculate the AR of a Cell. We first obtain the moment of inertia, **I**, in a frame of reference whose origin coincides with the center of mass of the cell, then diagonalize **I**, and calculate the AR, $r$, as the square root of the ratio of the two eigenvalues.

### S4. Procedure to calculate AR in our simulation

As stated in the main text, we characterize the cell shape via AR. To calculate the AR, we follow Ref. [17] and briefly discuss the method here. Consider one particular cell, represented by the set of points $\{x_i^1, x_i^2\}$, as schematically shown in Fig. S1. We first calculate the moment of inertia in a frame of reference whose origin coincides with the center of mass $(x_c^1, x_c^2)$ of the cell. The moment of inertia, **I**, is a $2 \times 2$ tensor in spatial dimension two. We next diagonalize **I** and obtain the two eigenvalues, $\Lambda_1$ and $\Lambda_2$. Without loss of generality, we assume that $\Lambda_1 \geq \Lambda_2$, and obtain the AR, $r = \sqrt{\Lambda_1/\Lambda_2}$. This process can be viewed as approximating the cell with an ellipse with the major and minor axes given by $\sqrt{\Lambda_1}$ and $\sqrt{\Lambda_2}$, respectively. For the CPM on the square lattice and the VM, we have written our own codes; for the *CompuCell3D* [12, 13], we have used `<Plugin Name="MomentOfInertia"/>` in the .XML file and using the example of `Demos/MomentOfInertia`, we have calculated the lengths of semiaxes in the `Steppables.py` file. We have obtained the AR of a cell by taking the ratio of lengths of semi-major and semi-minor axes.

### S5. Distribution of AR is independent of lattice type

In the main text, we have presented the results for the CPM on a square lattice and the VM, the latter being a continuum model. The agreements between the two models show that the results

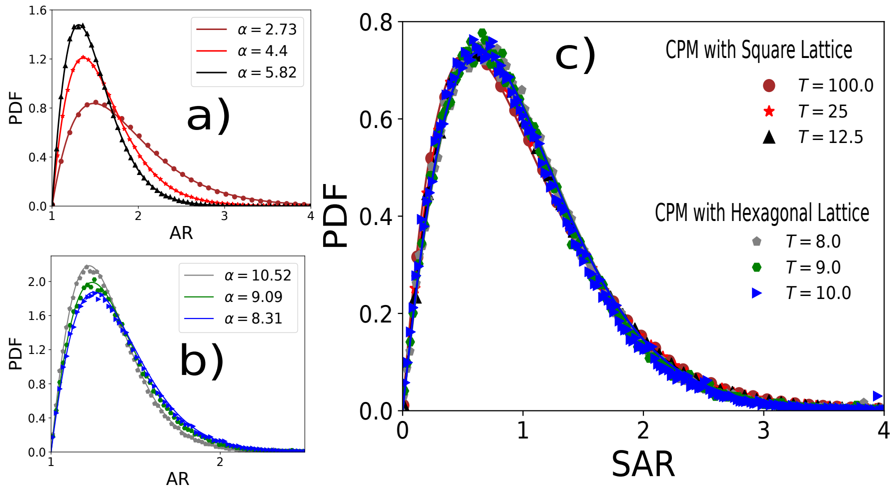


FIG. S2. Comparison among the CPM simulations on a square and hexagonal lattice. We have used $\lambda_A = 1.0$, $\lambda_P = 0.5$, $T$ values are quoted in (c). (a) PDF of $r$ on square lattice with $P_0 = 26$, (b) PDF of $r$ on hexagonal lattice with $P_0 = 25$. In both (a) and (b), symbols are simulation data, and lines represent the fits with Eq. (6) in the main text. Values of $\alpha$ are given in the figures. (c) PDFs for the $r_s$, for the same data as in (a) and (b), show a virtually universal behavior.

are independent of the lattice. As further evidence of this lattice independence, we present the results within the CPM on the square and the hexagonal lattices in Fig. S2. We show that our analytical theory agrees with simulations on both lattices.

Figures S2(a) and (b) show representative plots comparing the PDFs of $r$ within the CPM with the square and the hexagonal lattices, respectively. As discussed in the main text, our theory predicts almost universal behavior for the PDFs of $r_s$ (Fig. S2 c). This result shows that qualitative behaviors within the CPM are lattice-independent and agree with our analytical theory.

### S6. Distribution of AR does not depend on $\lambda_A$

One of the arguments within our mean-field theory is the following: since AR can be independent of $A$ and the shape should have the dominant contribution from the cortical properties, we can assume that individual cells satisfy the area constraint and ignore the area term in $\mathcal{H}$, Eq. (1) in the main text. We have shown that the area distribution is sharply peaked around the average area (Fig. 2(e) in the main text, and Fig. S5). We have also shown in the insets of Figs. 2(a) and (b) that $\alpha$ does not depend on $\lambda_A$, both within the CPM and the VM. We show in Fig. S3 that the PDFs of $r$ are almost overlapping with each other at different values of $\lambda_A$. Figure S3(c) shows the PDFs for $r_s$.

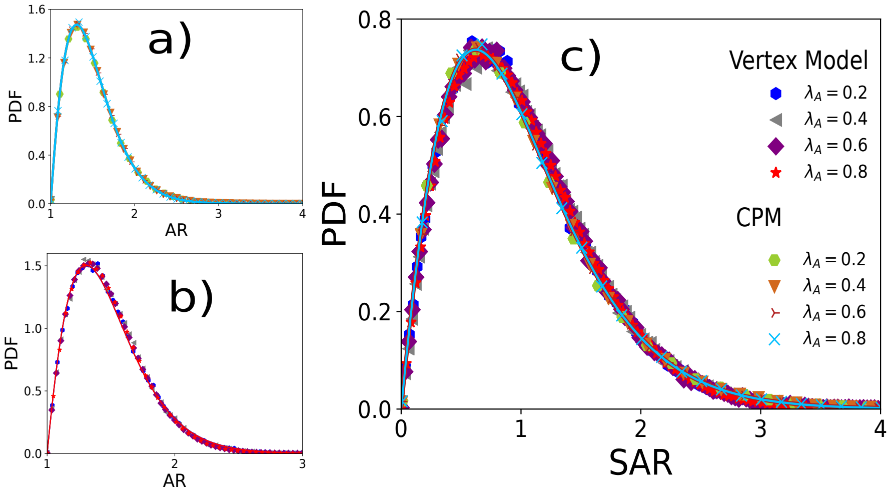


FIG. S3. PDFs of $r$ at different values of $\lambda_A$ within the CPM and the VM. Values of $\lambda_A$ are quoted in (c). (a) PDF of AR within the CPM for $\lambda_P = 0.5$ and $T = 12.5$ and (b) within the VM for $\lambda_P = 0.02$ and $T = 0.0025$ (c) PDFs of $r_s$, for the same data as in (a) and (b), show a virtually universal behavior. Symbols are simulation data, and lines represent the fits with Eq. (6) with $\alpha$ given in the main text in inset of Figs. 2(a) and (b).

### S7. Distribution of AR including cell division and apoptosis in the CPM simulation

We have ignored cell division and apoptosis in our theory since the rates of these processes are extremely low. However, even when these processes are significant, it is easy to see that their effects can only enter via $\alpha$. The reason behind this is that the other, algebraic part, comes from

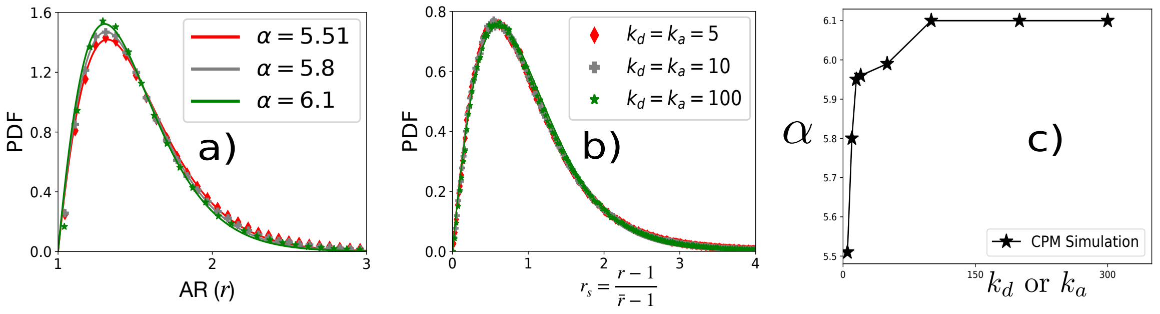


FIG. S4. PDFs of AR in the presence of cell division and apoptosis. (a) PDFs of AR at different values of $k_d = k_a$ (quoted in b) within CPM for $\lambda_P = 0.5$ and $T = 12.5$. (b) PDF of $r_s$ corresponding to the same data as in (a). (c) $\alpha$ vs $k_d$ or $k_a$ quickly reaches a constant with increasing $k_a$.

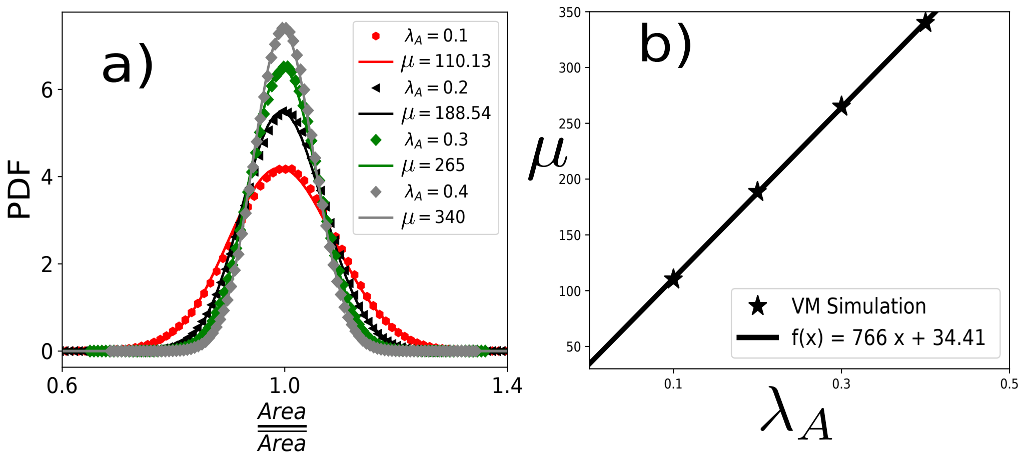


FIG. S5. Scaled area distribution within the VM. (a) PDF of the scaled area, $a$, within the VM at different $\lambda_A$ and fixed $P_0 = 3.7$, $\lambda_P = 0.02$, and $T = 0.0025$. Symbols are simulation data, and lines represent the fits with Eq. (7) of the main text. (b) $\mu$ linearly increases with $\lambda_A$.

the topology of the cell perimeter, that it is a closed-looped object; this property must remain the same even in the presence of these processes. Therefore, all the conclusions of our theory should remain valid even when the rates of division and apoptosis are significant.

To test this result, we have included cell division and apoptosis in our simulations. We keep the rate of division, $k_d^{-1}$, and apoptosis, $k_a^{-1}$, the same such that the average number of cells remains the same. Every $k_d$ time step, we randomly select a cell and divide it into two, with a randomly chosen division plane. To decide the division plane, we first calculate the center of mass (CoM) and then chose a point on the perimeter in a random direction. The line connecting the CoM and this point gives the division plane. The area of these two daughter cells then grows till it becomes of the order of $A_0$ (Eq. (1) in the main text). To avoid diving a cell that has just undergone division, we impose a cut-off on area ($\geq 32$) for selecting a cell for division. For apoptosis, every $k_a$ time step, we randomly choose a cell and assign it a target area $A_0 = 0$. We also relax the constraint that does not allow fragmentation in our simulation for the cells undergoing apoptosis.

We show the PDF of $r$ at three representative values of $k_a$ in Fig. S4(a), and the corresponding PDFs for $r_s$ are shown in Fig. S4(b). $k_a$ and $k_d$ are presented in units of the inverse of the Monte-Carlo time. We have fitted the PDFs of $r$ with Eq. (6) of the main text, and the values of $\alpha$ at different $k_a$ are shown in Fig. S4(c). We emphasize two observations: first, the PDFs of $r$ indeed fit well with Eq. (6) of the main text, and second, that $\alpha$ attains a constant value quite fast with increasing $k_a$, that is, decreasing the rate of division or apoptosis.

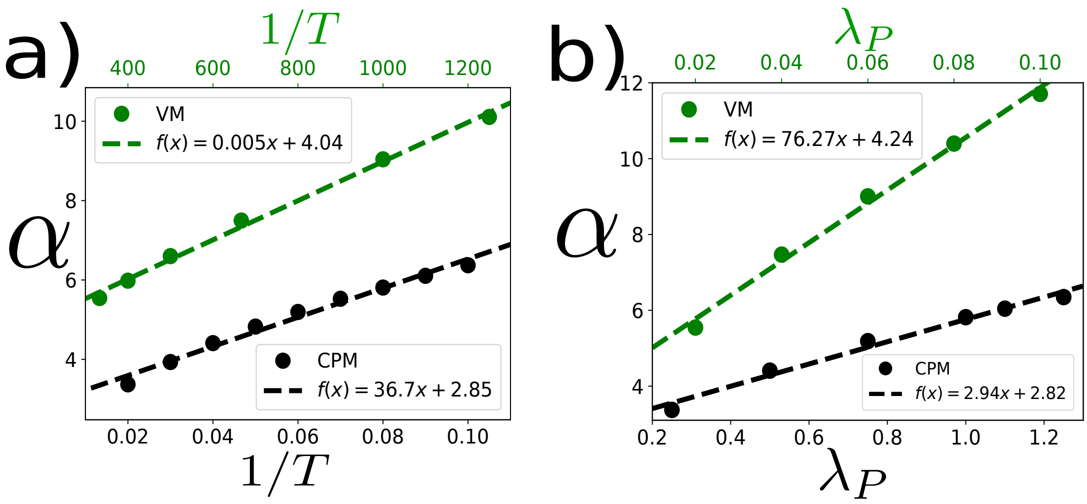


FIG. S6. Behavior of $\alpha$ with $T$ and $\lambda_P$. (a) $\alpha$ increases almost linearly as a function of $1/T$. Parameters for the simulations with the VM (represented by the green symbols and line) $\lambda_A = 0.5$, $\lambda_P = 0.02$, $P_0 = 3.7$. Within the CPM (black): $\lambda_A = 1.0$, $\lambda_P = 0.5$, and $P_0 = 26$. (b) $\alpha$ increases almost linearly as a function of $\lambda_P$. For the VM (green): $\lambda_A = 0.5$, $T = 0.003$, $P_0 = 3.7$. For the CPM (black): $\lambda_A = 1.0$, $T = 25$ ,$P_0 = 26$. Points represent simulation data and dashed lines represent the fits.

### S8. Scaled area distribution within the VM with varying $\lambda_A$

In the main text, we have shown the distribution of scaled area, $a$, for different values of $\lambda_A$ within CPM and their comparison with the analytical theory, Eq. (7). Here, in Fig. S5(a), we show the same distribution within the VM simulation and comparison with our theory. Figure S5(b) shows that $\mu$ linearly increases as $\lambda_A$ increases, similar to the result within the CPM (Fig. 2e, in the main text).

### S9. Dependence of $\alpha$ on $\lambda_P$ and $T$ within the CPM and the VM

In the main text, we have shown the linear dependence of $\alpha$ on $\lambda_P/T$: within both the CPM and the VM (Fig. 2d in the main text). Here, we show how $\alpha$ varies within both models when we change $\lambda_P$ and $T$, individually. Figure S6(a) shows the behavior of $\alpha$ as a function of $1/T$ and Fig. S6(b) as a function of $\lambda_P$: $\alpha$ increases almost linearly as $1/T$ or $\lambda_P$ increases, in agreement with our theory.

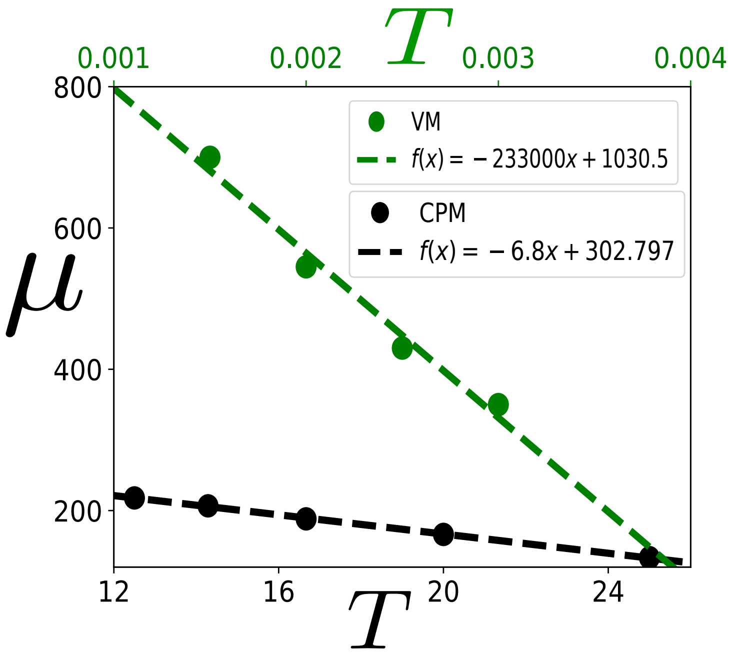


FIG. S7. The behavior of $\mu$ as a function of $T$. The coefficient $\mu$, in Eq. (7) in the main text, characterizes the PDF of scaled area, $a$. We show that $\mu$ decreases almost linearly as $T$ increases within both the CPM and the VM. In these simulations, the values of the parameters are as follows. For VM: $P_0 = 3.7$, $\lambda_A = 0.5$, and $\lambda_P = 0.02$; for CPM: $P_0 = 26$, $\lambda_A = 1.0$, and $\lambda_P = 0.5$.

### S10. Dependence of $\mu$ on $T$

Figure 2(f) in the main text shows that $\mu$ remains almost constant with varying $\lambda_P$. This behavior is expected since $\mu$ comes from the constraint of confluency and should be related to the area, whereas $\lambda_P$ is related to the perimeter constraint. On the other hand, from the same argument, we also expect $\mu$ to depend on $\lambda_A$; as shown in Fig. 2(e) in the main text, it increases linearly with $\lambda_A$. In addition, since $T$ controls the fluctuation in the system, we also expect $\mu$ to change with $T$; this dependence leads to different distributions of $a$ in epithelial systems. We show in Fig. S7 that $\mu$ almost linearly decreases as $T$ increases.

### S11. Fits with the HBEC data of AR

We have compared our theory for the AR, $r$, and the scaled AR, $r_s$, with the experimental data for three different systems: MDCK cells, asthmatic HBEC, and non-asthmatic HBEC cells. These data are taken from Ref. [17], where PDFs of $r$ and $r_s$ were presented as a function of maturation. As discussed in the main text, we have selected the data at three different times, and

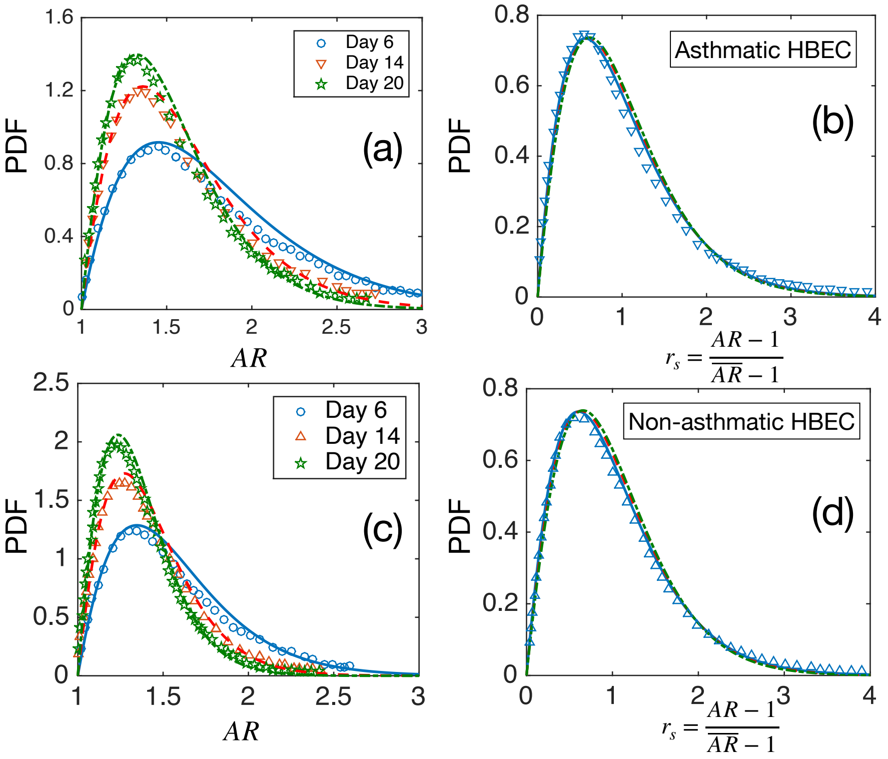


FIG. S8. Comparison of theory with experiments on HBEC cell monolayer. (a) Probability distribution function (PDF) of AR for asthmatic HBEC cells. Symbols are data taken from Ref. [17], and the lines represent fits with our theory, Eq. (6) in the main text. Values of $\alpha$ are shown in the main text, Table I. (b) PDF of the scaled AR, $r_s$ for the asthmatic HBEC cells. Lines are the theoretical predictions using the values of $\alpha$ obtained from the fits in (a), symbols represent one set of experimental data [17]. (c) and (d) The same as in (a) and (b), but for non-asthmatic HBEC cells.

the comparison for the MDCK cells are shown in Fig. 3(a) in the main text. Here, in Fig. S8, we present the comparisons for the asthmatic and non-asthmatic HBEC cells. We have collected only one set of experimental data for the PDF of $r_s$ in both the asthmatic and non-asthmatic systems, as all the data nearly overlap. The experimental data from different papers are collected using the software "WebPlotDigitizer" [18] that allows reading off the data from a digital plot.

[6] Wolfram Research Inc., "Mathematica, Version 12.0," Champaign, IL, 2019.

[18] A. Rohtagi, "Webplotdigitizer: Version 4.4," (2020).
